# Long-term memory reorganization of navigational episodes

**DOI:** 10.1101/2025.04.01.646566

**Authors:** Deetje Iggena, Thereza Schmelter, Patrizia M. Maier, Khaled Requieg, Carsten Finke, Kristian Hildebrand, Christoph J. Ploner

## Abstract

During navigation, the brain builds representations of self-motion and of environmental information for future action. The classic view suggests that these representations consolidate and eventually stabilize. However, there is no data on their fate at extended memory delays. Here, we investigated memory of navigational episodes from a real-world setting across memory delays of up to three decades. We show that memory of navigational episodes does not achieve a stable state, but rather continues to transform for many years. Our data suggest that at any given point in time, memory of navigational episodes is a changing combination of episode-independent schematic information with several interacting spatial representations that directly relate to a navigational episode and that show distinct decay rates. Consistent with recent accounts of memory reorganization, we further show that neither current theories of systems consolidation nor classic models of forgetting fully explain spatial memory performance at extended memory delays.

## 1. Introduction

When we navigate through an environment, we form representations of spatial information in memory. This information includes the paths we travel, the landmarks we encounter and the spatial relationships between these landmarks and ourselves ^1–4^. These spatial memory representations facilitate future behavior in previously visited locations, e.g. to find food or shelter. Everyday experience suggests that navigational episodes are rarely remembered in their entirety over extended periods. Instead, some details fade while other aspects remain salient, thus suggesting that long-term memory of navigational episodes may shift from a detailed cognitive map to a more gist-like spatial representation ^5^. However, although the transformative nature of memory consolidation has been stressed repeatedly ^6–9^, only few studies have investigated transformation of human spatial memory across real-life delays so far ^5,10^.

For natural environments, memory of spatial navigational episodes mostly occurs against the backdrop of semantic knowledge and patterns extraced from previous experiences that shape our expectations and guide actual behavior ^5,11^. For example, when we navigate a city, we expect the townhall to be in the centre of the city; when we go grocery shopping, we expect the commodities of a supermarket to be organized in semantic clusters; when we go hiking, we expect a valley behind a mountain etc. During individual navigational episodes, these spatial reference templates (schema) are combined with navigation-related information from a first-person perspective (egocentric) and a bird’s-eye perspective (allocentric). Studies suggest that memory of these representational modes relies on at least partially dissociable neural substrates ^5,12,13^.

There is currently sparse evidence whether the relative contributions of egocentric and allocentric representations and their substrates change with extended memory delays and how they interact with previous knowledge. For example, an fMRI study on mental navigation in large scale environments showed that involvement of the hippocampus decreases within a year when an environment is highly familiar and is navigated frequently after initial acquisition ^14^. A further study found that involvement of the hippocampus and retrosplenial cortex during a memory-guided navigation task depended on whether the environment was familiar for at least two years or newly learned ^15^. Conversely, a recent study suggests that the relative contributions of hippocampus and extrahippocampal regions to remote spatial memories are less dependent on the length of the memory delay, but rather on task demands such as the necessity to make precise spatial judgements ^16^. Moreover, for other memory domains, previous research found changes in memory representations that were between two weeks and two years old, but no changes for hippocampal and cortical representations of memories that were two or twelve years old, thus suggesting that memory consolidation and transformation may be limited to a few years after encoding ^17^.

Here, we investigated memory of real-world navigational episodes across extended memory delays. Former visitors of a zoo in Berlin were tested for their memory of the spatial layout of the zoo by using immersive virtual reality (VR) and 2D-maps. The delay between participants’ last navigational episode in the zoo and testing varied between six days and three decades, thus providing a unique opportunity to investigate memory delays that go far beyond previous studies of spatial memory consolidation.

Visitor performance was compared to zoo-naïve control subjects. We investigated whether egocentric and allocentric spatial memory representations continue to transform over extended time periods, whether their transformation can be explained with classic power functions of forgetting and how they interact with previous semantic knowledge.

## 2. Methods

### 2.1 Participants

A total of 134 participants were tested (104 zoo visitors, 30 zoo-naïve controls; 102 female and 32 male). All participants were recruited via online advertisement and on-site recruitment at the Charité-Universitätsmedizin Berlin. The first recruitment phase took place in August and September 2020 (58 participants) and the second recruitment phase from February 2022 to May 2023 (76 participants). Participants were between 14 and 71 years old, fluent in German, had normal or corrected-to-normal vision, normal hearing, reported being in good health and denied neuropsychiatric disorders or substance abuse. Participants were divided into two groups depending on whether they had visited the zoo at least once (zoo visitors vs. zoo-naïve controls). The number of zoo visits varied as well as the age at the last zoo visit (Table 1). All participants gave written informed consent. All experimental procedures were conducted in accordance with the Declaration of Helsinki and were approved by the local ethics committee of Charité-Universitätsmedizin Berlin.

**Table 1.** Descriptive participant data. Data are presented as percentage or median and IQR. Note: n.a. = not applicable.

|  | Zoo visitors (*N* = 104) | Zoo-naïve controls (*N* = 30) |
| --- | --- | --- |
| **Demographic** |  |  |
| Female/male (%) | 78/ 22 | 73/ 27 |
| Age (years) | 34 (26 – 51) | 27 (25 – 31) |
| Years of education | 17 (16 – 19) | 18.5 (17 – 19) |
| Right-/left-/binary handedness (%) | 95/ 4/ 1 | 73/ 20/ 7 |
| **Zoo visits** |  |  |
| Number of zoo visits | 4 (1.5 - 10) | n.a. |
| Time since first visit (days) | 6387 (2555 – 11406) | n.a. |
| Time since last visit (days) | 975 (183 – 3665) | n.a.. |
| Age at last visit (years) | 31 (21 – 40) | n.a.. |

### 2.2 Behavioral assessment

#### 2.2.1. The Berlin Zoo Task

Participants were tested on their spatial memory of the *Tierpark Berlin* which is one of two zoos in Berlin, Germany. The zoo is a 160-hectare site (Figure 1) and was founded in 1955. It contains several distinctive landmarks such as a castle, animal enclosures, zoo houses or restaurants that are easily distinguishable from each other.

**Figure 1.**
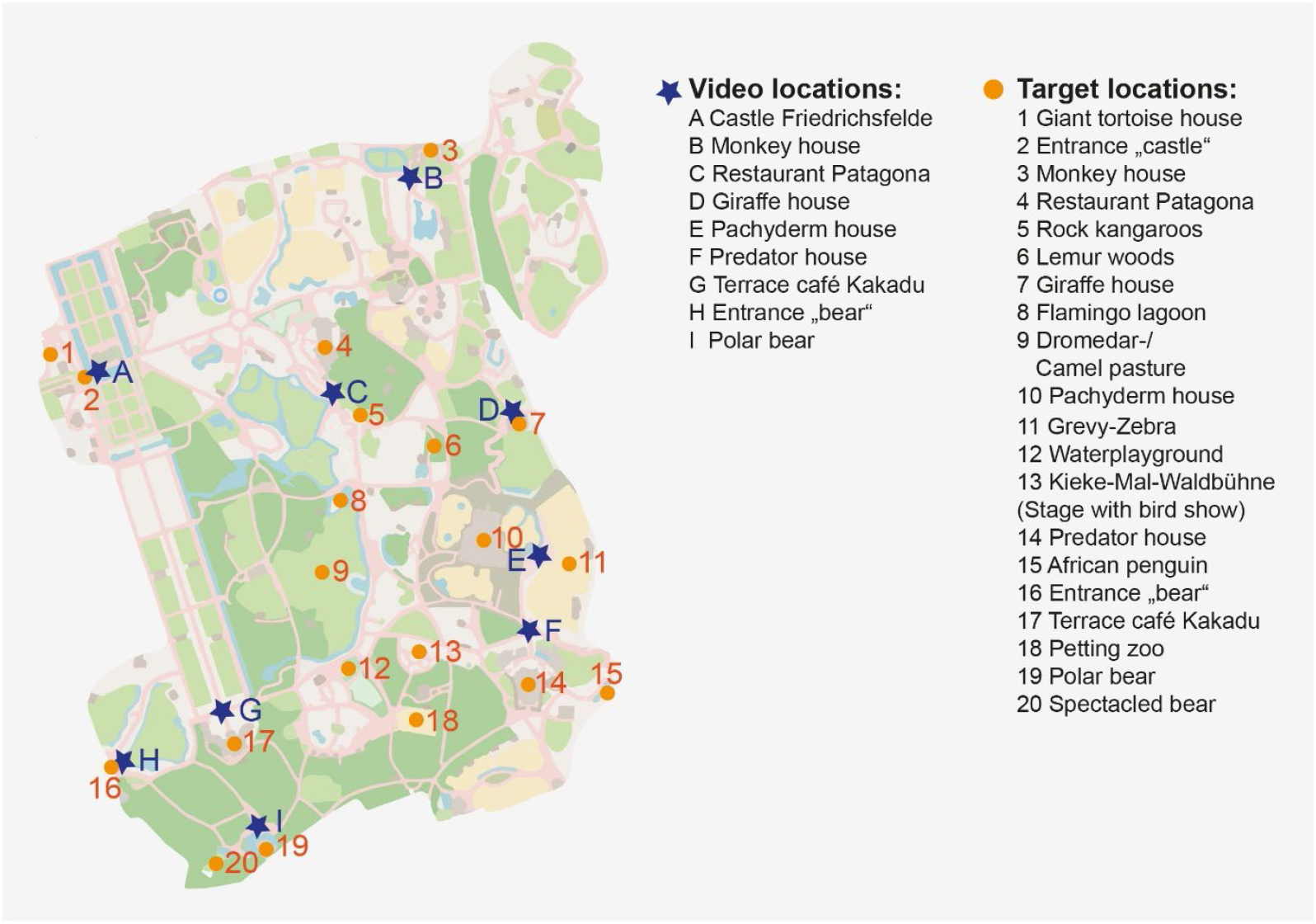
Overview map of the zoo - *Tierpark Berlin*. The locations of the video recordings are marked with a dark blue asterisk. The locations that served as targets are marked with an orange dot. Map data from OpenStreetMap.

The participants’ spatial memory of the zoo was tested with three different tasks assessing egocentric and allocentric spatial memory representations (Figure 2). In addition, participants completed questionnaires to determine covariates (see below). Individual data points per participant were excluded if a landmark that did not exist at the time of the last visit to the zoo was included in the experiment, e.g. the monkey house was only opened in 2000.

**Figure 2.**
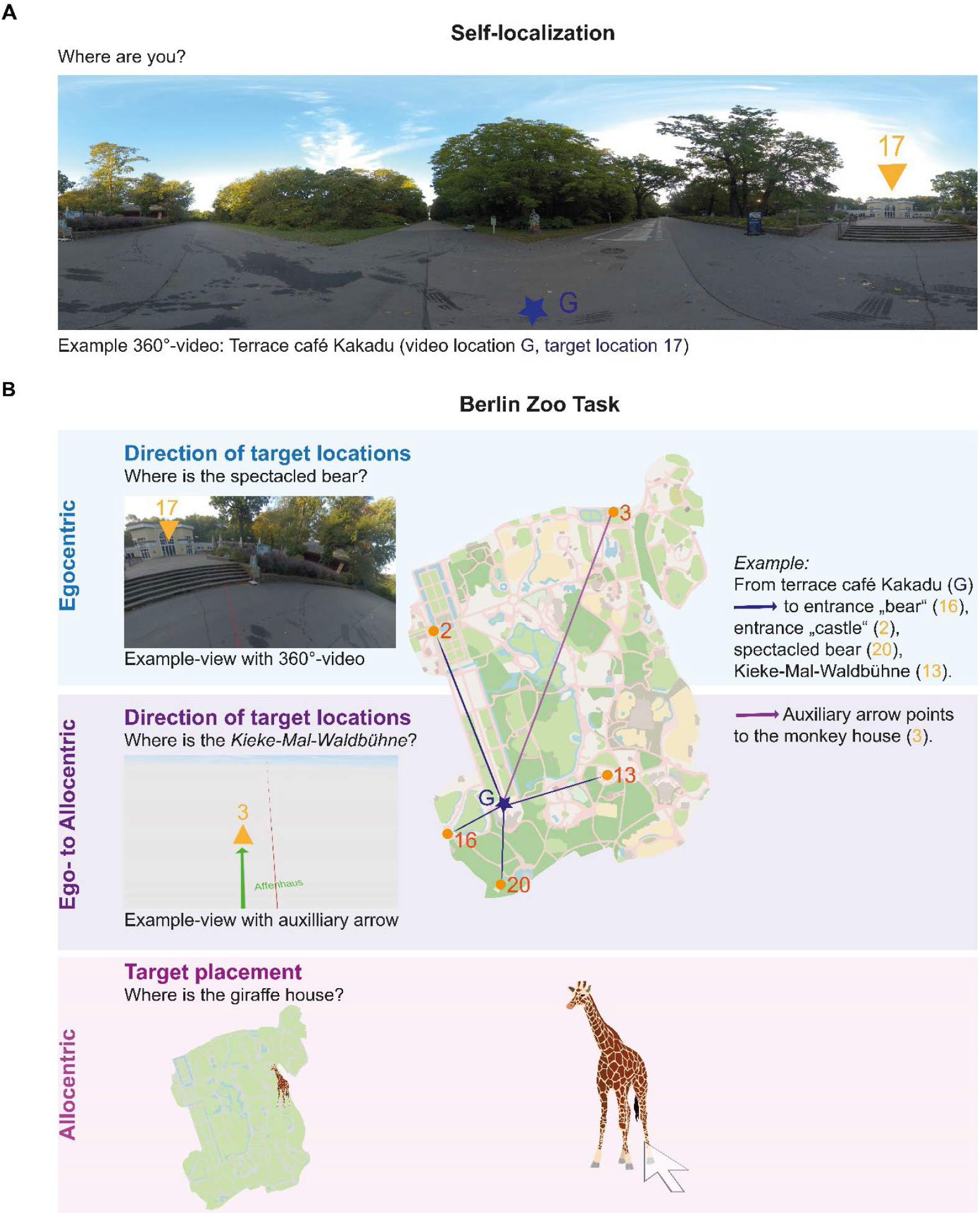
Experimental design. **A**, Self-localization. Participants are asked to locate themselves in the zoo using the environmental information provided in the 360° video; **B**, The Berlin Zoo task. The task consists of three different tasks that depend on egocentric and allocentric spatial representations of the zoo. Two tasks are performed from a first-person perspective in a virtual environment: Direction of target location asks participants to point in the direction of a specific location either by using the environmental information provided in the 360° video (egocentric); or by using an arrow pointing in the direction of a third location (egocentric to allocentric). A third task is performed from a bird’s eye view on a 2D screen. Target placement asks participants to place a specific location in the correct position on a mute map of the zoo (allocentric). Map data from OpenStreetMap.

#### 2.2.2. Experimental setup

For memory-guided pointing tasks from a first-person perspective, a total of nine 360° videos (ultra-high resolution, down sampled from 8k to 4k) were recorded at the zoo in October 2019. The 360° videos were recorded at different locations without overlapping features but at least one distinct environmental landmark (Figure 1 & 2A). The task was developed in Unity3D (version 2018.2.14f, Unity Technologies) and presented using an HTC VIVE Pro Eye virtual reality headset, provided by a mobile VR laboratory, to fully immerse participants in the zoo environment ^18^. The 360°-videos were mapped to the correct video locations using GPS coordinates and landmarks on a technical map of the zoo embedded in a virtual environment. In addition, the center of each target position was marked on the technical map. The technical map accurately reflected the spatial structure of the zoo and was provided by *Tierpark Berlin* authorities for research purposes.

The 2D map used for tasks from a bird’s eye view were programmed in MATLAB (Mathworks, R2018b) and presented with Psychtoolbox ^19^. The task was presented on a Lenovo Thinkpad X1 Carbon laptop (14.0-inch screen). The mute map used was extracted from Google Maps in January 2020 (map data: ©2020, Geo-Basis DE/BKG, (©2009), Google).

#### 2.2.3. Experimental procedure

Before testing, all participants were trained for the general design of the pointing tasks. Using a virtual reality headset, they were immersed in a 360° environment close to Brandenburg Gate - a well-known Berlin landmark - where they had to localize themselves and then point in the direction of several remote Berlin landmarks and to the north (supplementary material 1.1, table 1). Then, we assessed the accuracy of memories of navigational episodes in the zoo from a first-person perspective. For this purpose, participants were placed in immersive 360° videos of the zoo. Participants first had to localize themselves within the zoo and were then requested to point precisely to several different remote locations of the zoo (Figure 2, supplementary material 1.2 & 1.3, table 2 & 3). Afterwards, we assessed the accuracy of memories of navigational episodes in the zoo from a bird’s eye view by placing target locations on a 2D-map.

**Table 2.**
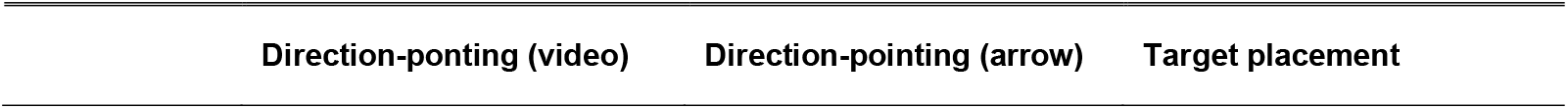
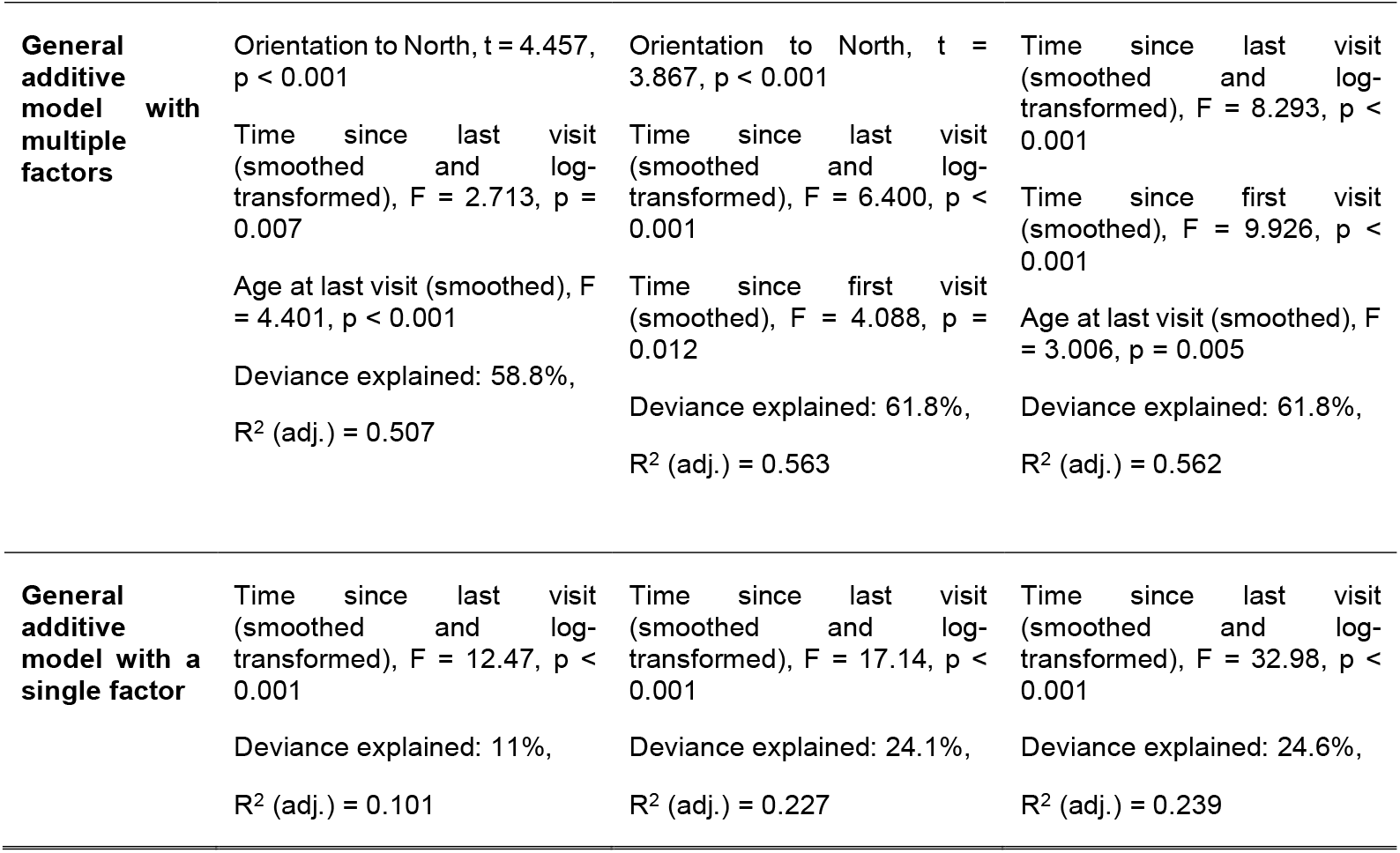
Table of results. Best fitting generalized additive model per task.

| Direction-ponting (video) | Direction-pointing (arrow) | Target placement |
| --- | --- | --- |

|  |  |  |  |
| --- | --- | --- | --- |
| **General additive model with multiple factors** | Orientation to North, $t = 4.457$, $p < 0.001$ | Orientation to North, $t = 3.867$, $p < 0.001$ | Time since last visit (smoothed and log-transformed), $F = 8.293$, $p < 0.001$ |
| | Time since last visit (smoothed and log-transformed), $F = 2.713$, $p = 0.007$ | Time since last visit (smoothed and log-transformed), $F = 6.400$, $p < 0.001$ | Time since first visit (smoothed), $F = 9.926$, $p < 0.001$ |
| | Age at last visit (smoothed), $F = 4.401$, $p < 0.001$ | Time since first visit (smoothed), $F = 4.088$, $p = 0.012$ | Age at last visit (smoothed), $F = 3.006$, $p = 0.005$ |
| | Deviance explained: 58.8%,<br>$R^2$ (adj.) = 0.507 | Deviance explained: 61.8%,<br>$R^2$ (adj.) = 0.563 | Deviance explained: 61.8%,<br>$R^2$ (adj.) = 0.562 |
| **General additive model with a single factor** | Time since last visit (smoothed and log-transformed), $F = 12.47$, $p < 0.001$ | Time since last visit (smoothed and log-transformed), $F = 17.14$, $p < 0.001$ | Time since last visit (smoothed and log-transformed), $F = 32.98$, $p < 0.001$ |
| | Deviance explained: 11%,<br>$R^2$ (adj.) = 0.101 | Deviance explained: 24.1%,<br>$R^2$ (adj.) = 0.227 | Deviance explained: 24.6%,<br>$R^2$ (adj.) = 0.239 |

##### Self-localization

Participants were asked to report on their location within the zoo verbally (Figure 2A). The answer was rated as either correct or incorrect. If the answer was incorrect or the participants could not localize themselves at all, they were told the correct answer to avoid errors in the subsequent pointing tasks. A total of nine video locations were tested (Figure 1 & 2, supplementary material 1.3, table 3). To ensure that the locations were recognizable, all locations contained at least one specific environmental landmark, such as the terrace in front of Cafe Kakadu (Figure 2A). Each self-localization was followed by two pointing tasks (Figure 2B, supplementary material 1.3 & 1.4, table 3, figure 1).

##### Direction pointing with video (egocentric)

In this pointing task, participants were immersed in the 360°-video and asked to point in the direction of several specific target locations of the zoo, for which they had to relate knowledge of their own position to the features of their immediate environment using the 360°-video. The pointing task always began with pointing to the entrances of the zoo, because each participants’ navigational episodes must have started by entering the zoo via one of these locations. In addition, they were asked to point to two other locations of the zoo thus yielding a total of three to four location trials per video location (supplementary material 1.3, table 3). Target locations were always separated by at least 45° of visual angle to induce orienting movements of participants. These additional target locations appeared only once during the task and were not repeated at subsequent video positions. In addition, all participants were asked to point to the north at the end of each pointing task to test their global spatial orientation. After completion of the pointing task in the 360° video environment, the next video location was shown and participants were again asked to localize themselves, followed by the next pointing task. The order of the video locations was randomized, but the target locations and their order were kept constant for each video location (supplementary material 1.3, table 3). A total of 20 target locations were tested (Figure 1).

For the analysis of pointing performance, the angular deviation from the target location was standardized and compared to the z-score per query of the zoo-naïve controls. This procedure was chosen to account for logical inferences about possible target locations in zoo-naïve participants, e.g., signs or alleys guiding to landmarks.

##### Direction pointing with arrow (egocentric to allocentric)

In this pointing task, the 360° video was removed, and the participants found themselves in a neutral virtual environment with a white floor and a blue sky with several clouds. No landmarks for spatial orientation were provided. They were now asked to imagine that they were standing in the place they had immersed themselves in during self– localization. As an orientation aid, a green circle with a green arrow was projected onto the white floor, pointing in the direction of a second location, e.g. the monkey house (Figure 2B, supplementary material 1.3, table 3). The participants were asked to turn in the direction of the arrow, followed by pointing in the direction of several distinct locations (same locations and order as in direction-pointing with video). This task required the participants to integrate knowledge of their own position and the position of two locations into a global map and to relate these locations to each other spatially. After the completion of the pointing task in the neutral environment, the environment faded out and the original video was shown again (supplementary material 1.4, figure 1).

For analysis of pointing performance, the angular deviation from the target location was standardized and compared to the z-score per query of the zoo-naïve controls as in the pointing-task with video.

##### Target placement (allocentric)

In this task, participants were asked to place target locations on a 2D-map of the zoo from a bird’s eye view (Figure 2B). This target placement task requires allocentric representations of the zoo. To place the target locations, participants first saw an isolated image of the location, followed by a fixation cross for 1 second, then the map appeared, and the specific location had to be placed on the map by moving the mouse and confirming with a mouse click. Participants had 20 seconds to decide where to place the specific location and were explicitly asked to place the target locations as precisely as possible even if they could not remember the exact location. After the decision, a fixation cross was displayed again for 1 second and the next trial started. The same 20 target locations were tested as for the pointing tasks (Figure 1, supplementary material 1.3, table 3).

For the analysis of placement performance, the distance between the chosen location and the target location was measured in pixels, standardized and compared to the z-score per query of the zoo-naïve controls. Trials in which objects were placed outside the silent map were excluded from the analysis.

The order of the three tasks was fixed to minimize the risk of transfer of spatial knowledge between tasks (supplementary material 1.4, figure 1).

### 2.4. Assessment of covariates

All participants completed socio-demographic questionnaires. These included age, biological sex, years of education, and handedness. Additionally, all zoo visitors stated how often they had visited the zoo in total and provided the approximate dates of their first and last visits. The age at the last zoo visit was calculated using the age at testing and the date of the last zoo visit (Table 1).

To assess spatial abilities, all participants completed a German version of the Santa Barbara Sense of Direction Scale (SBSOD) ^20^. The questionnaire was rated on a Likert scale from 1 to 7. For the SBSODs, a higher final score indicated a better sense of direction.

### 2.5. Statistical analysis

Statistical analyses were performed in RStudio (v. 4.2). To compare the responses of the zoo visitors with the performance of the control participants and to account for the different localization properties in the three tasks, we first converted the raw scores of all participants into z-scores. These z-scores are based on the average performance of the control participants for each query in the pointing task and for each target placement in the target placement task. We then aggregated the data per participant in each task.

Next, we visually analyzed the z-scores of the three tasks and examined skewness and kurtosis of the data. Then, we used the Shapiro-Wilk test to formally check whether the data of the three tasks were normally distributed. Since the assumption of a normal distribution had to be partially rejected, we compared zoo visitors and the zoo-naïve controls with a non-parametric one-sided Mann-Whitney U Test (supplementary material 1.5, table 4). Experimental tasks were compared using a Kruskal-Wallis-Test. Post-hoc tests were conducted using pairwise Mann-Whitney U tests. P values were corrected for multiple comparisons using the Bonferroni method. For effect sizes, *r* was calculated for the Mann-Whitney U test (small: 0.1 <= r < 0.3; medium: 0.3 <= r < 0.5; large: r >= 0.5) and η^2^ was calculated for the Kruskal-Wallis-test (small: 0.01 <= η^2^ < 0.06; medium: 0.06 <= η^2^ < 0.14; large: η^2^ >= 0.14)

Correlations between task performance and covariates were calculated using Spearman’s rank sum correlation. Comparison of the correlation-coefficients between tasks was conducted with a Fisher’s z-test.

To uncover the relationship between time and spatial memory performance, we employed eight different regression models to investigate which regression would best explain the collected data (figure 3, supplementary material 1.6, table 5). For logarithmic, exponential and power regression, the respective data were log-transformed after being converted to a positive value. Due to a sample size of n = 104, we performed regression validation with leave-one-out cross-validation (LOOCV). The Akaike Information Criterion (AIC) was chosen for model selection as it is also valid for non-linear regressions. Further model descriptors were the mean square error (MSE), the root mean square error (RMSE), R^2^.adjusted and the Bayesian Information Criterion (BIC) (supplementary material 1.6, table 5). A two-way ANOVA was performed for each regression analysis to examine whether spatial memory performance changed over time across the tasks. Post-hoc pairwise comparisons of slopes were performed and p values were corrected for multiple comparisons using the Tukey method. Effect sizes were assessed using ꞷ^2^ (small: 0.01 <= ꞷ^2^ < 0.06; medium: 0.06 <= ꞷ^2^ < 0.14; large: ꞷ^2^ >= 0.14).

**Figure 3.**
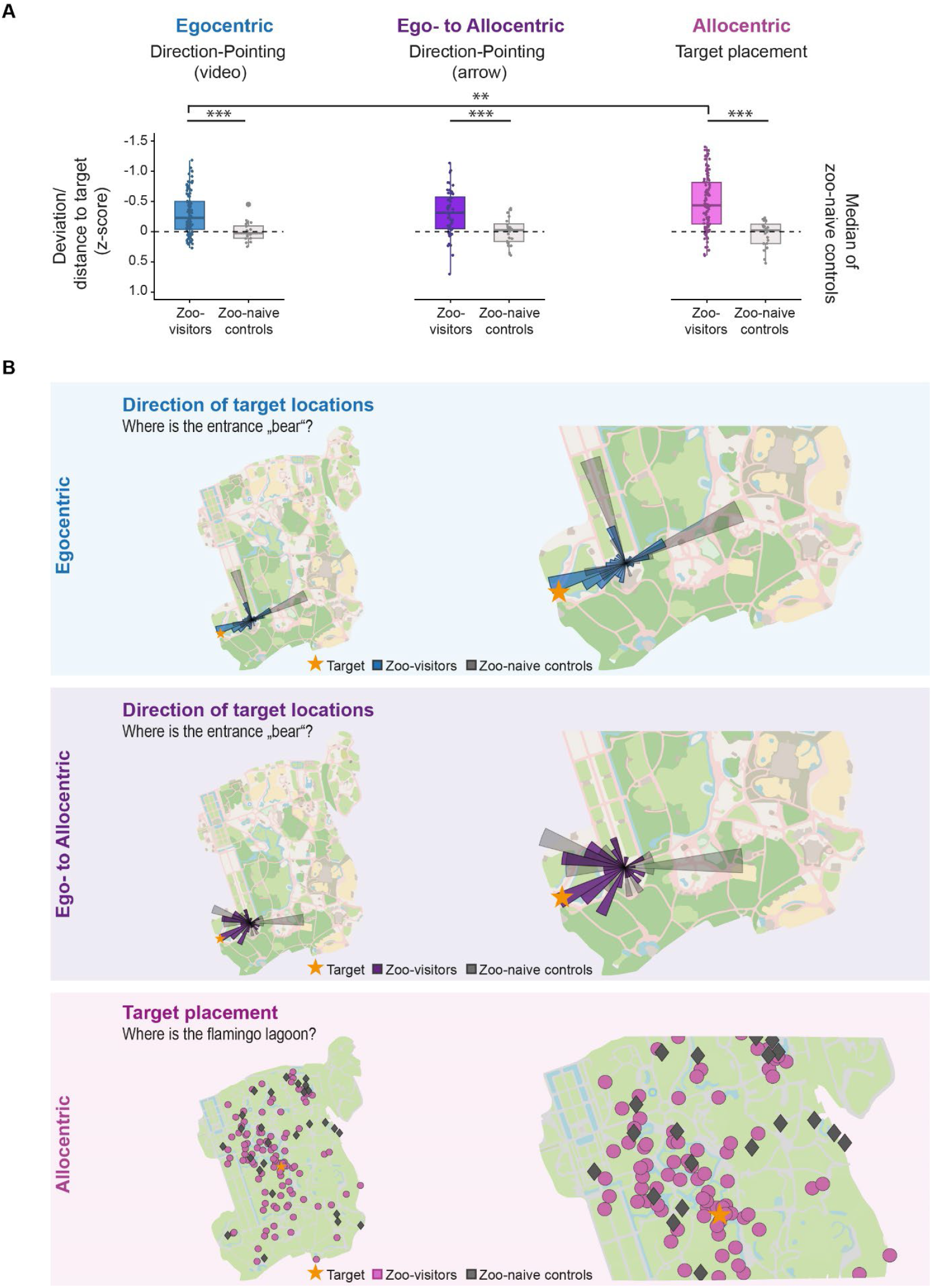
Spatial memory performance of zoo visitors compared to zoo-naïve controls. **A)** Zoo visitors showed a lower deviation and distance (z-score) from target locations and outperformed zoo-naïve controls in all three tasks. In zoo visitors, overall performance was better for target placement than for pointing into the direction of target locations. p < 0.05 = *; p < 0.01 = **; p < 0.001 = ***. **B)** Example maps for spatial memory performance in the three tasks. Top: Pointing in the direction of the target location from the video location using a 360° video; middle: Pointing in the direction of the target location from the imagined video location using an arrow; bottom: Target placement on a silent map of the zoo. Coloured direction indicators and dots show responses of zoo visitors, gray direction indicators and rhombs show responses of zoo-naïve controls. Map data from OpenStreetMap.

To model the forgetting rate, the equation *-1*.*84/(log10(time)*^*1*.*25*^ *+ 1*.*84* was used to simulate the “Ebbinghaus curve” as the classic and highly influential mathematical model ^21,22^. For comparison with the respective tasks of our study, the equation was adjusted by multiplication with the calculated highest assumed z-score of each task, which was extracted from the best fitting regression equation (Direction-pointing (video), -1.757; Direction-pointing (arrow), -1.934; target placement, -2.249; supplementary material 1.7, table 6). Additionally, only corresponding time points were chosen to simulate the Ebbinghaus curve.

To investigate the contribution of multiple factors to spatial memory performance, we applied a generalized additive model (GAM). In addition to the time since last visit, we examined other zoo-visit related factors, including the time since the first visit, the total number of zoo visits, and age at the last visit. Personal characteristics such as age and educational level were also considered. Additionally, spatial strategies and abilities, such as northward orientation – the ability to point to the north -, were included in the model. The final model selected was the simplest model that best explained spatial memory consolidation, incorporating only the most relevant and significant contributing factors (Table 2).

The level of significance was set to the conventional level of *p* < 0.05.

### 2.6. Data availability

The data of this study is accessible at OSF

## 3. Results

### 3.1 Performance in the Berlin Zoo Task depends on memory of real-world navigational episodes

We first examined whether performance in the Berlin Zoo task depends on spatial representations acquired during navigation at the zoo and whether previous semantic knowledge contributes to performance. We therefore tested whether zoo visitors recognized video locations, whether zoo visitors outperform zoo-naïve controls on all three tasks of the Berlin Zoo task and whether performance of zoo-naïve participants is better than chance.

In 77% of cases (IQR: 67 – 89%), zoo visitors were able to correctly localize themselves using the 360°-video (supplementary material 1.2, table 2). The rate of correct responses varied between 17% and 100% across locations. We then compared zoo visitors’ performance with that of zoo-naïve controls (Figure 3A & B, supplementary material 1.5, table 5). For all three tasks, we found that the average performance of zoo visitors was better than that of zoo-naïve controls (direction-pointing (video), W = 771, p < 0.001, r = 0.377; direction-pointing (arrow), W = 257, p < 0.001, r = 0.430; target placement, W = 576, p < 0.001, r = 0.464). Comparison of performance across all three tasks revealed that memory performance in the target placement task was better compared to direction-pointing with video support (χ^2^ = 11.09, p = 0.004, η^2^ = 0.029; direction-pointing (video) vs. direction-pointing (arrow), p = 1.0; direction-pointing (video) vs. target placement, p = 0.004; direction-pointing (arrow) vs. target placement, p = 0.097). These results indicate that performance in the Berlin Zoo task was neither random nor solely influenced by environmental stimuli but rather depended on the availability of a representation of preceding navigational episodes in the zoo.

We next analyzed whether performance in the Berlin Zoo task was additionally driven by environmental information that supports target localization independently of navigational episodes in the zoo. We thus compared performance of zoo-naïve controls with random performance in all three tasks (direction-pointing (video): 90°; direction-pointing (arrow): 90°; target placement: 0.5). We found that the performance of zoo-naïve controls in all tasks was better than chance (direction-pointing (video), p < 0.001, r = 0.836; direction-pointing (arrow), p < 0.001, r = 0.701; target placement, p < 0.001, r = 0.873). These results suggest that environmental information additionally activated spatial semantic knowledge that may have facilitated inferences about target locations.

### 3.2 Memory of real-world navigational episodes transforms non-linearly across decades

Visual inspection of the z-scores indicated a non-linear relationship between time and spatial memory performance for all three tasks (Figure 4A). To describe the relationship further, we correlated spatial memory performance with the time elapsed since the last visit to the zoo. For all three tasks, we found that spatial memory performance continuously decreased with increasing time since last visit (Figure 4A, ssupplementary material 1.6, table 5; direction-pointing (video), Rho = 0.25, p = 0.010; direction-pointing (arrow), Rho = 0.370, p = 0.004; target placement, Rho = 0.380, p < 0.001). When comparing performance between tasks, we found no differences in correlations (direction-pointing (video) vs. direction-pointing (arrow), z = -1.521, p = 0.128; direction-pointing (video) vs. target placement, z = - 1.306, p = 0.192; direction-pointing (arrow) vs. target placement, z = 0.175, p = 0.861).

**Figure 4.**
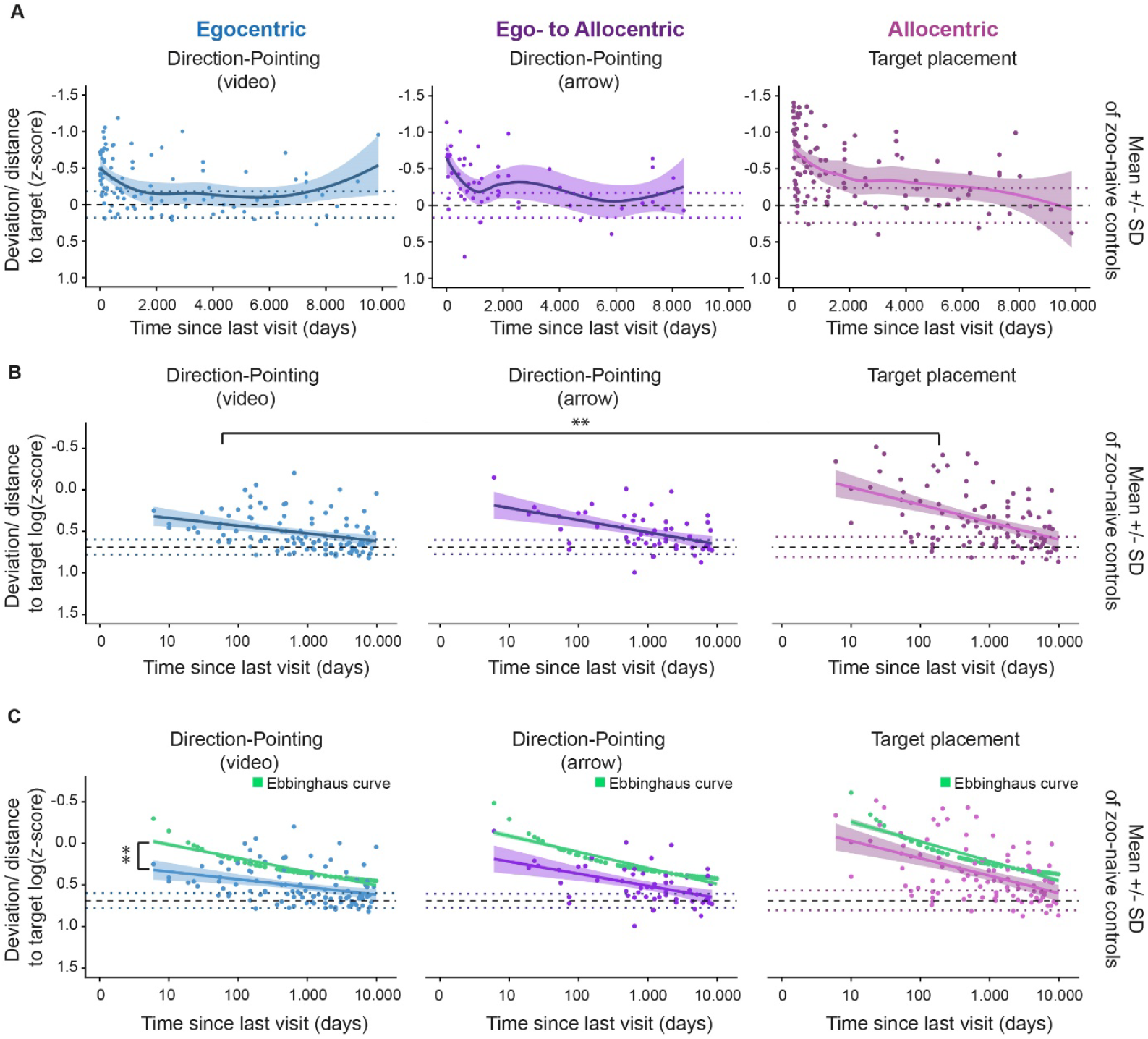
Memory of navigational episodes in the zoo over a period of six days to three decades since last visit. **A)** For zoo visitors, data of all three tasks was locally weighted and smoothed for visual inspection. Dotted lines indicate the first standard deviation above and below the mean performance of zoo-naïve controls. Dashed line indicates mean performance of zoo-naïve controls. Performance on all tasks was significantly correlated with the time elapsed since the last visit to the zoo (direction-pointing (video), Rho = 0.25, p = 0.010; direction-pointing (arrow), Rho = 0.370, p = 0.004; target placement, Rho = 0.380, p < 0.001). **B)** The power regression for all three tasks revealed a decline in memory performance as a function of the time since the last visit to the zoo. Dotted lines indicate the first standard deviation above and below the mean performance of zoo-naïve controls – were shifted into the positive range before applying a log transformation. Dashed line indicates mean performance of zoo-naïve controls. The regression slope differed significantly between direction-pointing (video) and target placement (F_(2,256)_ = 3.735, p = 0.025, ꞷ^2^ = 0.02). p < 0.05 = *; p < 0.01 = **; p < 0.001 = ***. **C)** The power regressions of all tasks are compared with the Ebbinghaus equation and the simulated data, starting with the highest possible score according to the power regressions. Dotted lines indicate the first upper and lower standard deviation of the zoo-naïve controls – z-values were transformed in a positive value for log-transformation. On all tasks zoo visitors performed less efficiently than predicted by the Ebbinghaus curve. A difference was found between the slopes for the direction-pointing with video task and the simulated data (F_(2,54)_ = 4.736, p = 0.013, ꞷ^2^ = 0.11). p < 0.05 = *; p < 0.01 = **; p < 0.001 = ***. Dashed lines: mean of zoo-naïve controls; dotted lines: upper and lower standard deviations of zoo-naïve controls. Note that z-scores were calculated for each individual query in the tasks. The data presented here is aggregated per participant. Mean and SD refer to these aggregated values per participant.

To investigate the causal relationship between time and spatial memory performance further, we performed a regression analysis of spatial memory performance for all three tasks over time. After employing eight different regression models, the Akaike information criterion (AIC) revealed that a power regression best explained the impact of time since last visit on spatial memory performance in all three tasks (supplementary material 1.6, table 5). Comparison of the slopes of the regression lines also showed a significant interaction between time elapsed since last visit and task, thus suggesting that memory decay differed between direction-pointing with video and target placement, i.e. between the purely egocentric and purely allocentric tasks (F_(2,256)_ = 3.735, p = 0.025, ꞷ^2^ = 0.02; direction-pointing (video) vs. direction-pointing (arrow), p = 0.568; direction-pointing (video) vs. target-placement, p = 0.019; direction-pointing (arrow) vs. target placement, p = 0.447).

We then analyzed at which point in time spatial memory performance of zoo visitors corresponded to that of zoo-naïve controls, i.e. at which point in time performance was no longer mainly driven by representations of past navigational episodes, but rather by semantic knowledge and logical inference. The most appropriate and interpretable model was chosen for the calculation, which was a power regression for all tasks. For direction-pointing with video, the average performance of the zoo visitors reached the mean performance of the zoo-naïve controls after about 52,560 days (144 years [upper SD reached by 5,658 days (15.5 years)]; supplementary material 1.7, table 6). For the ‘direction-pointing with arrow’ task, average performance of the zoo visitors reached performance of the zoo-naïve controls after 14,965 days (41 years [upper SD reached by 3,945 days (10.8 years)]). In the ‘target placement’ task, we found a performance overlap at 24,090 days (66 years [upper SD reached by 6,300 days (17.3 years)]).

Since performance of zoo visitors did not stabilize but rather continued to decrease to the level of zoo-naïve controls, the relationship between time and spatial memory performance in our study can equally be conceived in frameworks of consolidation and of forgetting. We therefore investigated whether representations of past navigational episodes can be modelled with a classic logarithmic function of forgetting such as the highly influential Ebbinghaus curve (Ebbinghaus 1885, Murre & Dros 2015, Della Salla et al. 2024). After aligning the curves with the same starting value on a hypothetical first day for each task, we compared the curves and found an interaction effect for direction-pointing (video) between time and curve for the experimentally collected datasets and the hypothetical Ebbinghaus datasets (Figure 4C F_(1, 202)_ = 7.128, p = 0.008, ꞷ^2^ = 0.03). This suggests that the time course of egocentric spatial memories differs significantly from the Ebbinghaus curve. By contrast, for direction-pointing (arrow) and target placement, we found a difference between the experimental and hypothetical data sets and a time effect, but no interaction effect. This suggests a time course similar to the Ebbinghaus curve for allocentric spatial representations (direction-pointing (arrow): Interaction, F_(1,155)_ = 2. 881, p = 0.092, ꞷ^2^ = 0.01, experimental vs. hypothetical data, F_(1,155)_ = 107.615, p < 0.001, ꞷ^2^ = 0.40, Time, F_(1,155)_ = 197.109, p < 0.001, ꞷ^2^ = 0. 55; target placement: F(1,_202)_ = 1.495, p = 0.223, ꞷ^2^ = 0.002, experimental vs. hypothetical data, F_(1,202)_ = 37.818, p < 0.001, ꞷ^2^ = 0.15, time, F_(1,202)_ = 148.453, p < 0.001, ꞷ^2^ = 0.42).

### 3.3 Memory of real-world navigational episodes also depends on time-independent factors

Further analysis revealed that the time since last visit only explained 11 to 25% of the observed variance in spatial memory performance (Direction-pointing (video): R^2^ (adj.) = 0.101, deviance explained = 11%; Direction-pointing (arrow): R^2^ (adj.) = 0.227, deviance explained = 24.1%; Target placement: R^2^ (adj.) = 0.239, deviance explained = 24.6%).

We therefore analyzed whether additional factors in conjunction with time since the last visit could better explain spatial memory performance of zoo visitors in the Berlin Zoo task. For the direction-pointing with video task, the best generalized additive model (GAM) explained 58.8% of the variance, incorporating time since the last visit, northward orientation during the task, and age at the time of the last visit. For the direction-pointing with arrow task, the optimal GAM explained 61.8% of the variance, with contributing factors including time since the last visit, northward orientation, and time since the first visit to the zoo. Similarly, for the target placement task, the best GAM accounted for 61.8% of the variance, with time since the last visit, time since the first visit, and age at the last visit as the most relevant predictors. The final model presented in Table 2 is the simplest model that best explains spatial memory performance, incorporating only the most significant contributing factors.

## 4. Discussion

We investigated memory of navigational episodes from a real-world setting, a zoo in Berlin, across memory delays of up to three decades. Contrary to what the concept of memory consolidation implies, our results show that zoo visitors’ memory of their navigational experience does not achieve a stable state but rather continues to transform for many years. Our data is consistent with the hypothesis that at any given point in time, memory of navigational episodes is a continuously changing combination of episode-related egocentric and allocentric spatial representations with episode-independent schematic knowledge. These representations show distinct decay rates that do not necessarily follow classic models of forgetting and suggest distinct but interacting neural mechanisms of maintenance even years after a navigational experience.

Historically, the term consolidation suggests the transition of memory representations from an initial labile to a more stable state. Although initially applied to memory experiments on a timescale of minutes, the term was later also used to explain temporally graded amnesia in patients with temporal lobe damage ^9,23^. The seeming preservation of remote autobiographical memories in these patients was compatible with the hypothesis that consolidation at the level of brain systems is a process that takes years and ultimately renders memories resistant even against brain damage. It further suggested that consolidation leads to a progressive shift from hippocampal to extrahippocampal brain regions for storage of memories, a hypothesis that is still under debate ^6,8,9,24^. For obvious methodological reasons, it has proven difficult to investigate the time course of systems consolidation experimentally. Therefore, only few studies have addressed the hypothetical years-long process in human participants without brain damage. In a series of fMRI experiments on normal human subjects recalling autobiographical episodes from up to ten or twelve years ago, no differences were found between memories that were two or twelve years old, thus suggesting that autobiographical memory consolidation may be largely complete by, at most, two years after memory formation ^17,25^. Several fMRI studies also investigated consolidation of spatial memory by comparing navigational performance or distance judgements in environments learned years ago. However, it has remained equivocal whether consolidation-related changes in brain activation are mainly driven by time-related factors or the demands of the task ^14,16,26^. The question of whether spatial memory consolidation indeed levels off at some point in time therefore still needs experimental evaluation.

Our retrospective study design spanned a period of time that exceeds previous prospective and retrospective longitudinal memory experiments. The variance in time since the last zoo visit allowed for a detailed analysis of the time-dependency of performance in all three memory tasks. We observed that the time course of all studied spatial memory representations was non-linear with a loss of accuracy especially during the initial 1000 – 2000 days. However, performance continued to change at longer memory delays, suggesting that spatial representations continued to transform even at delays that are otherwise considered to be a domain of autobiographical memory testing. The resulting non-linear forgetting curve for all spatial representations in our study resembles power functions from classical memory experiments such as the Ebbinghaus experiments ^21,22,27^. Although primarily obtained with non-sense verbal material on a time-scale of minutes to months, studies suggest the applicability to other memory domains, including visual memory ^28,29^. However, a comparison of the hypothetical data for spatial representations calculated by the Ebbinghaus equation with our real data revealed that overall, a stronger memory decay of the real data was observed and that the time course of egocentric memory representations was less steep than predicted by the equation.

The distinct time-dependency of memory performance between tasks further suggests that memories of initial navigational episodes are not stored as holistic representations that consolidate or deteriorate in an all-or-none mode. Rather, increasing the memory delay seems to lead to a continuous shift in the respective contributions of different spatial representations to overall spatial memory performance, reflected in distinct forgetting curves for the three tasks. One factor that critically determines the slope of memory retention across time is intermittent retrieval ^27^. Egocentric representations, as assessed with our video-assisted pointing task, are thus in a privileged position, as they re-create the initial navigational experience from a first-person perspective rather than an abstracted allocentric map. Studies have repeatedly shown that egocentric perspective has a beneficial effect on subsequent memory performance ^30–32^. Spontaneous intermittent retrieval of a navigational episode between last visit and testing may thus have occurred from a first-person perspective and may have affected the slope of memory retention of egocentric representations, but not or much lesser of allocentric representations. Egocentric representations may thus be conceived as the main carriers of episodic information following a navigational experience. In line with this hypothesis, our analyses suggest that the average performance of zoo visitors in the egocentric task would reach the mean performance of zoo-naïve controls after 144 years, i.e. clearly after the average human lifetime. Conversely, although allocentric representations, as assessed by the target placement task, also relate to initial navigational episodes, their lack of detail and birds-eye perspective render allocentric representations more to gist-like summaries of the spatial layout of the navigational episodes rather than to representations that allow for intermittent retrieval of navigational experiences. They may thus be conceived as abstracted templates that may ultimately contribute to an overall spatial “zoo”-schema that is independent of a distinct navigational episode ^5^.

The behavior of zoo-naïve control subjects indeed shows that performance was not solely driven by memory of navigational episodes in the Berlin zoo but also by a schematic representation independent from the zoo. Thus, visual stimuli may have guided behavior by prompting retrieval of schematic representations of previously visited environments that share spatial features with the Berlin zoo ^5^. Supporting this notion, we found that zoo-naïve control subjects performed above chance level in all three tasks. These findings therefore suggest that memories of navigational episodes acquired in real-world settings are not purely episodic in nature. Instead, they emerge as a combination of the actual navigational experience and semantic knowledge that facilitates logical inferences and decision-making in environments with overlapping spatial features. In this framework, cognitive maps, spatial gist and spatial schema are not three successive steps of abstraction of an initial experience, but rather three classes of spatial representations with distinct relationships to navigational episodes that exist in parallel and mutually influence each other ^5^.

It should, however, be pointed out that time since last visit explained only up to a quarter of the variance in memory performance across the three tasks. Instead, a combination of factors such as time since last visit, age at last visit, and spatial orientation skills can explain up to two-thirds of the variance in memory performance. Time thus appears as only one of many factors that determines the interplay of spatial representations. This finding aligns with the notion that spatial navigation is influenced by several factors such as age and individual spatial abilities ^12,13,33–35^.

Taken together, our findings suggest that memory of navigational episodes emerges from the interaction of parallel representations, each likely supported by distinct neural substrates within a widespread hippocampal-neocortical network ^5,11,34^. These substrates are dynamically activated to varying degrees based on past experiences and current task demands, with each activation influencing the organization and structure of the retrieved memory. We assume that this process continuously reshapes memory of navigational episodes, reflecting the brain’s ability to adapt and restructure memory networks in response to changing cognitive demands and environmental contexts. This process can be described most appropriately as a memory reorganization process, as it seems likely that consolidation and forgetting of spatial memories never ends ^6,8,9,36^. Combining retrieval of remote navigational episodes with fMRI measurements may ultimately reveal the underlying neural network dynamics.

## Open science practices

Data is available at osf

## Acknowledgements

We thank the study participants for their kind support. We thank Anton Moritz John for technical assistance. We thank all employees of the Berlin Zoo – *Tierpark Berlin* for their kind support, especially Anna Ohl and Katharina Sperling. Copyright information for Open Map Data is available at openstreetmap.org/copyright.

