## Supplement for "Long-term memory reorganization of navigational episodes"

### Supplementary material

### 1.1

**Table 1.** Descriptive location data for training task. Pointing from video location to target location.

| Trial | Video location | Target location | Euclidean distance | Video to target location angle | Auxiliary arrow points to |
|---|---|---|---|---|---|
| **Direction of target locations (ego- to allocentric)** | | | | | |
| 1 | Pariser Platz | Brandenburg Gate | 68m | -89.6° | TV Tower |
| 2 | Pariser Platz | Parliament | 886m | -38.6° | TV Tower |
| 3 | Pariser Platz | North | n.a. | 0° | TV Tower |
| **Direction of target locations (egocentric)** | | | | | |
| 1 | Pariser Platz | Brandenburg Gate | 702m | -89.6° | n.a. |
| 2 | Pariser Platz | Parliament | 886m | -38.6° | n.a. |
| 3 | Pariser Platz | North | n.a. | 0° | n.a. |

### 1.2

**Table 2.** Descriptive target location data. Data presented as median and IQR or percentage of correct responses. Data on self-localization only available for video locations.

| Location no. | Location name | **Familiarity** (Score 1 – 7) | **Self-Localization** (correct first response in %) |
|---|---|---|---|
| 1 | Giant tortoise house | 2.0 (2.0 – 5.0) | n.a. |
| 2 | Castle Friedrichsfelde | 4.0 (2.0 – 6.0) | 90% |
| 3 | Monkey house | 3.0 (2.0 – 4.0) | 17% |
| 4 | Restaurant Patagona | 3.0 (1.0 – 4.0) | 65% |
| 5 | Rock kangaroos | 3.0 (2.0 – 4.0) | n.a. |
| 6 | Lemur woods | 2.0 (1.0 – 4.0) | n.a. |
| 7 | Giraffe house | 4.0 (3.0 – 6.0) | 100% |
| 8 | Flamingo lagoon | 3.0 (2.0 – 5.0) | n.a. |
| 9 | Dromedar-/ Camel pasture | 4.0 (2.0 – 5.0) | n.a. |

| | | | |
|---|---|---|---|
| 10 | Pachyderm house | 4.5 (2.8 – 6.0) | 99% |
| 11 | Grevy-Zebra | 3.0 (2.0 – 4.0) | n.a. |
| 12 | Water playground | 2.0 (1.0 – 4.0) | n.a, |
| 13 | Kieke-Mal-Waldbühne (bird show) | 1.0 (1.0 – 2.0) | n.a. |
| 14 | Predator house | 4.0 (2.3 – 6.0) | 79% |
| 15 | African penguin | 3.0 (2.0 – 5.0) | n.a. |
| 16 | Entrance (Bear) | 6.0 (4.0 – 7.0) | 94% |
| 17 | Terrace café Kakadu | 3.0 (2.0 – 5.0) | 95% |
| 18 | Petting zoo | 4.0 (2.0 – 5.0) | n.a. |
| 19 | Polar bear | 5.0 (3.0 – 6.0) | 70% |
| 20 | Spectacled bear | 4.0 (2.0 – 5.0) | n.a. |

## 1.3

**Table 3.** Descriptive location data. Pointing from video location to target location. Letters and numbers refer to the overview map provided in figure 1. The order of video locations was randomized, the trial order was fixed.

| Trial | Video location | Target location | Euclidean distance | Video to target location angle | Auxiliary arrow points to |
|---|---|---|---|---|---|
| | **1. Direction of target locations (ego- to allocentric)** | | | | |
| 1 | A Castle Friedrichsfelde | 16 Entrance (Bear) | 702m | 176.6° | Predator house |
| 2 | A Castle Friedrichsfelde | 11 Grevy-Zebra | 886m | 115.5° | Predator house |
| 3 | A Castle Friedrichsfelde | 17 Terrace café Kakadu | 692m | 158.3° | Predator house |
| 4 | A Castle Friedrichsfelde | North | n.a. | 0.0° | Predator house |
| | **2. Direction of target locations (egocentric)** | | | | |
| 1 | A Castle Friedrichsfelde | 16 Entrance (Bear) | 702m | 176.6° | n.a. |
| 2 | A Castle Friedrichsfelde | 11 Grevy-Zebra | 886m | 115.5° | n.a. |
| 3 | A Castle Friedrichsfelde | 17 Terrace café Kakadu | 692m | 158.3° | n.a. |
| 4 | A Castle Friedrichsfelde | North | n.a. | 0.0° | n.a. |

| | | | | | |
|---|---|---|---|---|---|
| | **1. Direction of target locations (ego- to allocentric)** | | | | |
| 1 | B Monkey house | 2 Entry (Castle) | 659m | - 121.5° | Restaurant Patagona |
| 2 | B Monkey house | 16 Entry (Bear) | 920m | - 153.1° | Restaurant Patagona |
| 3 | B Monkey house | 7 Giraffe house | 508m | 155.3° | Restaurant Patagona |
| 4 | B Monkey house | 6 Lemur woods | 540m | 174.3° | Restaurant Patagona |
| 5 | B Monkey house | North | n.a. | 0.0° | Restaurant Patagona |
| | **2. Direction of target locations (egocentric)** | | | | |
| 1 | B Monkey house | 2 Entry (Castle) | 659m | - 121.5° | n.a. |
| 2 | B Monkey house | 16 Entry (Bear) | 920m | - 153.1° | n.a. |
| 3 | B Monkey house | 7 Giraffe house | 508m | 155.3° | n.a. |
| 4 | B Monkey house | 6 Lemur woods | 540m | 174.3° | n.a. |
| 5 | B Monkey house | North | n.a. | 0.0° | n.a. |
| | **1. Direction of target locations (ego- to allocentric)** | | | | |
| 1 | C Restaurant Patagona | 16 Entry (Bear) | 814m | - 150.4° | Giraffe house |
| 2 | C Restaurant Patagona | 2 Entry (Castle) | 364m | - 86.5° | Giraffe house |
| 3 | C Restaurant Patagona | 12 Water playground | 571m | 173.7° | Giraffe house |
| 4 | C Restaurant Patagona | 3 Monkey house | 390m | 25.1° | Giraffe house |
| 5 | C Restaurant Patagona | North | n.a. | 0.0° | Giraffe house |
| | **2. Direction of target locations (egocentric)** | | | | |
| 1 | C Restaurant Patagona | 16 Entry (Bear) | 814m | - 150.4° | n.a. |
| 2 | C Restaurant Patagona | 2 Entry (Castle) | 364m | - 86.5° | n.a. |
| 3 | C Restaurant Patagona | 12 Water playground | 571m | 173.7° | n.a. |
| 4 | C Restaurant Patagona | 3 Monkey house | 390m | 25.1° | n.a. |
| 5 | C Restaurant Patagona | North | n.a. | 0.0° | n.a. |
| | **1. Direction of target locations (ego- to allocentric)** | | | | |
| 1 | D Giraffe house | 16 Entry (Bear) | 920m | - 131.1° | Terrace café Kakadu |
| 2 | D Giraffe house | 2 Entry (Castle) | 711m | - 85.8° | Terrace café Kakadu |

| | | | | | |
|---|---|---|---|---|---|
| 3 | D Giraffe house | 9 Dromedar-/ Camel pasture | 427m | - 129.5° | Terrace café Kakadu |
| 4 | D Giraffe house | 5 Rock kangaroos | 281m | - 91.0° | Terrace café Kakadu |
| 5 | D Giraffe house | North | n.a. | 0.0° | Terrace café Kakadu |

**2. Direction of target locations (egocentric)**

| | | | | | |
|---|---|---|---|---|---|
| 1 | D Giraffe house | 16 Entry (Bear) | 920m | - 131.1° | n.a. |
| 2 | D Giraffe house | 2 Entry (Castle) | 711m | - 85.8° | n.a. |
| 3 | D Giraffe house | 9 Dromedar-/ Camel pasture | 427m | - 129.5° | n.a. |
| 4 | D Giraffe house | 5 Rock kangaroos | 281m | - 91.0° | n.a. |
| 5 | D Giraffe house | North | n.a. | 0.0° | n.a. |

**1. Direction of target locations (ego- to allocentric)**

| | | | | | |
|---|---|---|---|---|---|
| 1 | E Pachyderm house | 2 Entry (Castle) | 716m | - 71.9 | Monkey house |
| 2 | E Pachyderm house | 16 Entry (Bear) | 751m | - 117.2° | Monkey house |
| 3 | E Pachyderm house | 15 African penguin | 327m | 162.1° | Monkey house |
| 4 | E Pachyderm house | 19 Polar bear | 646m | - 137.0 | Monkey house |
| 5 | E Pachyderm house | North | n.a. | 0.0° | Monkey house |

**2. Direction of target locations (egocentric)**

| | | | | | |
|---|---|---|---|---|---|
| 1 | E Pachyderm house | 2 Entry (Castle) | 716m | - 71.9 | n.a. |
| 2 | E Pachyderm house | 16 Entry (Bear) | 751m | - 117.2° | n.a. |
| 3 | E Pachyderm house | 15 African penguin | 327m | 162.1° | n.a. |
| 4 | E Pachyderm house | 19 Polar bear | 646m | - 137.0 | n.a. |
| 5 | E Pachyderm house | North | n.a. | 0.0° | n.a. |

**1. Direction of target locations (ego- to allocentric)**

| | | | | | |
|---|---|---|---|---|---|
| 1 | F Predator house | 2 Entry (Castle) | 915m | - 60.9° | Polar bear |
| 2 | F Predator house | 16 Entry (Bear) | 746m | - 107.0° | Polar bear |
| 3 | F Predator house | 10 Pachyderm house | 251m | - 28.7° | Polar bear |
| 4 | F Predator house | 18 Petting Zoo | 207m | - 127.2° | Polar bear |
| 5 | F Predator house | North | n.a. | 0.0° | Polar bear |

| | | | | | |
|---|---|---|---|---|---|
| | **2. Direction of target locations (egocentric)** | | | | |
| 1 | F Predator house | 2 Entry (Castle) | 915m | - 60.9° | n.a. |
| 2 | F Predator house | 16 Entry (Bear) | 746m | - 107.0° | n.a. |
| 3 | F Predator house | 10 Pachyderm house | 251m | - 28.7° | n.a. |
| 4 | F Predator house | 18 Petting Zoo | 207m | - 127.2° | n.a. |
| 5 | F Predator house | North | n.a. | 0.0° | n.a. |
| | **1. Direction of target locations (ego- to allocentric)** | | | | |
| 1 | G Terrace café Kakadu | 16 Entry (Bear) | 221m | - 113.9° | Monkey house |
| 2 | G Terrace café Kakadu | 2 Entry (Castle) | 692m | - 23.6° | Monkey house |
| 3 | G Terrace café Kakadu | 20 Spectacled bear | 216m | - 177.1° | Monkey house |
| 4 | G Terrace café Kakadu | 13 Kieke-Mal-Waldbühne (bird show) | 245m | 75.2° | Monkey house |
| 5 | G Terrace café Kakadu | North | n.a. | 0.0° | Monkey house |
| | **2. Direction of target locations (egocentric)** | | | | |
| 1 | G Terrace café Kakadu | 16 Entry (Bear) | 221m | - 113.9° | n.a. |
| 2 | G Terrace café Kakadu | 2 Entry (Castle) | 692m | - 23.6° | n.a. |
| 3 | G Terrace café Kakadu | 20 Spectacled bear | 216m | - 177.1° | n.a. |
| 4 | G Terrace café Kakadu | 13 Kieke-Mal-Waldbühne (bird show) | 245m | 75.2° | n.a. |
| 5 | G Terrace café Kakadu | North | n.a. | 0.0° | n.a. |
| | **1. Direction of target locations (ego- to allocentric)** | | | | |
| 1 | H Entry (Bear) | 2 Entry (Castle) | 702m | - 5.8° | Polar bear |
| 2 | H Entry (Bear) | 8 Flamingo lagoon | 601m | 40.7° | Polar bear |
| 3 | H Entry (Bear) | 1 Giant tortoise | 713m | -9.5° | Polar bear |
| 4 | H Entry (Bear) | North | n.a. | 0.0° | Polar bear |
| | **2. Direction of target locations (egocentric)** | | | | |
| 1 | H Entry (Bear) | 2 Entry (Castle) | 702m | - 5.8° | n.a. |
| 2 | H Entry (Bear) | 8 Flamingo lagoon | 601m | 40.7° | n.a. |

| | | | | | |
|---|---|---|---|---|---|
| 3 | H Entry (Bear) | 1 Giant tortoise | 713m | -9.5° | n.a. |
| 4 | H Entry (Bear) | North | n.a. | 0.0° | n.a. |
| | **1. Direction of target locations (ego- to allocentric)** | | | | |
| 1 | I Polar bear | 16 Entry (Bear) | 304m | - 66.4 | Pachyderm house |
| 2 | I Polar bear | 2 Entry (Castle) | 885m | - 21.7° | Pachyderm house |
| 3 | I Polar bear | 14 Predator house | 444m | 62.6° | Pachyderm house |
| 4 | I Polar bear | 4 Restaurant Patagona | 874m | 8.1° | Pachyderm house |
| 5 | I Polar bear | North | n.a. | 0.0° | Pachyderm house |
| | **2. Direction of target locations (egocentric)** | | | | |
| 1 | I Polar bear | 16 Entry (Bear) | 304m | - 66.4 | n.a. |
| 2 | I Polar bear | 2 Entry (Castle) | 885m | - 21.7° | n.a. |
| 3 | I Polar bear | 14 Predator house | 444m | 62.6° | n.a. |
| 4 | I Polar bear | 4 Restaurant Patagona | 874m | 8.1° | n.a. |
| 5 | I Polar bear | North | n.a. | 0.0° | n.a. |

## 1.4

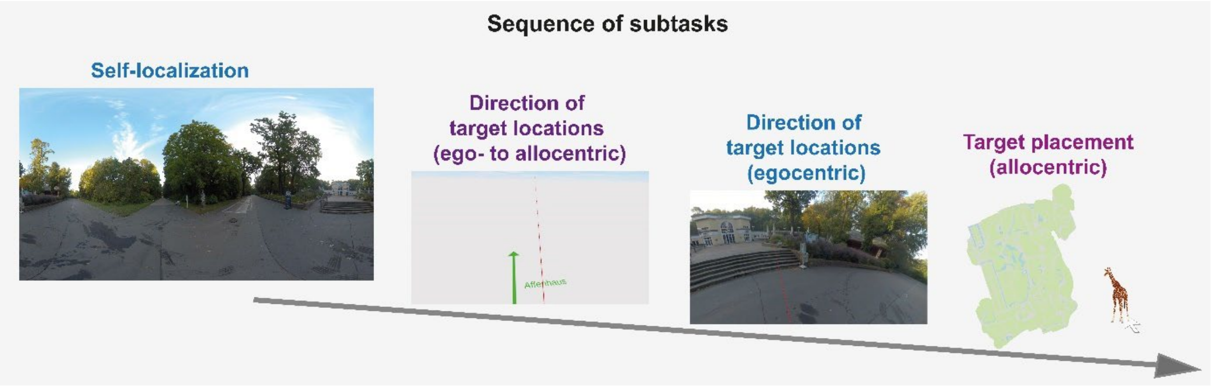


**Figure 1.** The order of the subtasks in the Berlin Zoo task. To avoid a transfer of spatial knowledge between the subtasks, self-localization was followed by the pointing task with arrow, followed by the pointing task with video and finally the target placement.

## 1.5

**Table 4.** Table of results for the comparison of memory accuracy as z-score between zoo visitors and zoo-naive control per subtask. Data presented as median and IQR.

| | **Zoo visitors** | **Zoo-naïve control** | ***p-value*** |
|---|---|---|---|
| **Self-localization (%)** | 77 (67 – 89) | 77 (67 – 77) | 0.014 |
| **Direction-Pointing (video)** | -0.226 (-0.498 - -0.036) | -0.016 (-0.143 - 0.138) | < 0.001 |
| **Direction-Pointing (arrow)** | -0.310 (-0.576 - -0.048) | -0.021 (-0.134 - 0.169) | < 0.001 |
| **Target placement** | -0.436 (-0.815 - -0.125) | -0.030 (-0.170 – 0.134) | < 0.001 |

## 1.6

**Table 5.** Table of results. Correlation and functions explaining the variance in spatial memory performance by elapsed time since last visit.

| | **Direction-pointing (video) a** | **Direction-pointing (arrow) b** | **Target placement c** | **Test statistic** |
|---|---|---|---|---|
| **Spearman's correlation** | Rho = 0.25,<br>p = 0.010 | Rho = 0.370,<br>p = 0.004 | Rho = 0.380,<br>$p < 0.001$ | a vs. b: z = -1.521, p = 0.128<br>a vs. c: z = -1.306, p = 0.192<br>b vs. c: z = -0.175, p = 0.861 |
| **Linear regression** | $R_{adj} = 0.061$<br>$F_{(1,100)} = 7.591$<br>p = 0.007<br>RSE = 0.332<br>MSE = 0.113<br>RMSE = 0.336<br>AIC = 68.340<br>BIC = 76.215 | $R_{adj} = 0.096$<br>$F_{(1,53)} = 6.717$<br>p = 0.012<br>RSE = 0.350<br>MSE = 0.125<br>RMSE = 0.354<br>AIC = 44.47<br>BIC = 40.49 | $R_{adj} = 0.171$<br>$F_{(1,100)} = 21.76$<br>$p < 0.001$<br>RSE = 0.407<br>MSE = 0.170<br>RMSE = 0.412<br>AIC = 110.24<br>BIC = 118.12 | Subtask:<br>$F_{(2,256)} = 8.418$, $p < 0.001$<br>Time since last visit:<br>$F_{(1,256)} = 34.311$, $p < 0.001$<br>Subtask*elapsed time:<br>$F_{(2,256)} = 1.801$, p = 0.166 |
| **Logarithmic regression**<br>(log-transformation of the predictor variable) | $R_{adj} = 0.129$<br>$F_{(1,100)} = 16$<br>$p < 0.001$<br>RSE = 0.138<br>MSE = 0.104<br>RMSE = 0.323<br>AIC = 103.02<br>BIC = 110.90 | $R_{adj} = 0.212$<br>$F_{(1,53)} = 15.55$<br>$p < 0.001$<br>RSE = 0.326<br>MSE = 0.110<br>RMSE = 0.332<br>AIC = 36.88<br>BIC = 42.90 | $R_{adj} = 0.238$<br>$F_{(1,100)} = 32.55$<br>$p < 0.001$<br>RSE = 0.390<br>MSE = 0.157<br>RMSE = 0.396<br>AIC = 101.58<br>BIC = 109.45 | Subtask:<br>$F_{(2,256)} = 9.162$, $p < 0.001$<br>Time since last visit:<br>$F_{(1,256)} = 60.326$, $p < 0.001$<br>Subtask*elapsed time:<br>$F_{(2,256)} = 1.780$, p = 0.171 |
| **Exponential regression** | $R_{adj} = 0.052$<br>$F_{(1,100)} = 6.546$ | $R_{adj} = 0.100$<br>$F_{(1,53)} = 6.977$ | $R_{adj} = 0.157$<br>$F_{(1,100)} = 19.76$ | Subtask:<br>$F_{(2,256)} = 10.181$, $p < 0.001$ |

| | | | | |
|---|---|---|---|---|
| (log-transformation of the dependent variable) | p = 0.012<br>RSE = 0.215<br>MSE = 0.761<br>RMSE = 0.872<br>AIC = -19.74<br>BIC = -11.86 | p = 0.011<br>RSE = 0.217<br>MSE = 0.781<br>RMSE = 0.884<br>AIC = -8.17<br>BIC = -2.15 | p < 0.001<br>RSE = 0.309<br>MSE = 0.895<br>RMSE = 0.946<br>AIC = 53.68<br>BIC = 61.55 | Time since last visit:<br>$F_{(1,256)}$ = 32.080, p < 0.001<br>Subtask*elapsed time:<br>$F_{(2,256)}$ = 2.605, p = 0.076 |
| **Power regression**<br>(log-transformation for the dependent and predictor variable) | $R_{adj}$ = 0.107<br>$F_{(1,100)}$ = 13.08<br>p < 0.001<br>RSE = 0.209<br>MSE = 0.753<br>RMSE = 0.868<br>AIC = -25.81<br>BIC = -17.93 | $R_{adj}$ = 0.237<br>$F_{(1,53)}$ = 17.77<br>p < 0.001<br>RSE = 0.120<br>MSE = 0.768<br>RMSE = 0.876<br>AIC = -17.27<br>BIC = -11.25 | $R_{adj}$ = 0.245<br>$F_{(1,100)}$ = 34.38<br>p < 0.001<br>RSE = 0.291<br>MSE = 0.882<br>RMSE = 0.939<br>AIC = 41.93<br>BIC = 49.80 | Subtask:<br>$F_{(2,256)}$ = 11.253, p < 0.001<br>Time since last visit:<br>$F_{(1,256)}$ = 60.680, p < 0.001<br>Subtask*elapsed time:<br>$F_{(2,256)}$ = 3.735, **p = 0.025**<br>Post-hoc comparison (Tukey):<br>a vs. b: p = 0.568<br>a vs. c: **p = 0.019**<br>b vs. c: p = 0.447 |
| **Quadratic polynomic regression** | $R_{adj}$ = 0.163<br>$F_{(2,99)}$ = 10.86<br>p < 0.001<br>RSE = 0.313<br>MSE = 0.103<br>RMSE = 0.321<br>AIC = 57.57<br>BIC = 68.07 | $R_{adj}$ = 0.153<br>$F_{(2,,52)}$ = 5.865<br>p = 0.005<br>RSE = 0.339<br>MSE = 0.121<br>RMSE = 0.348<br>AIC = 41.84<br>BIC = 49.87 | $R_{adj}$ = 0.181<br>$F_{(2,99)}$ = 12.19<br>p < 0.001<br>RSE = 0.404<br>MSE = 0.172<br>RMSE = 0.415<br>AIC = 109.87<br>BIC = 120.37 | Subtask:<br>$F_{(2,253)}$ = 8.7928, p < 0.001<br>Time since last visit:<br>$F_{(2,253)}$ = 24.442, p < 0.001<br>Subtask*elapsed time:<br>$F_{(4,253)}$ = 1.271, p = 0.282 |
| **Cubic polynomic regression** | $R_{adj}$ = 0.155<br>$F_{(3,98)}$ = 7.172<br>p < 0.001<br>RSE = 0.315<br>MSE = 0.110<br>RMSE = 0.332<br>AIC = 59.56<br>BIC = 72.68 | $R_{adj}$ = 0.139<br>$F_{(3,,51)}$ = 3.908<br>p = 0.014<br>RSE = 0.341<br>MSE = 0.125<br>RMSE = 0.354<br>AIC = 43.65<br>BIC = 53.68 | $R_{adj}$ = 0.209<br>$F_{(3,98)}$ = 9.867<br>p < 0.001<br>RSE = 0.404<br>MSE = 0.165<br>RMSE = 0.406<br>AIC = 107.40<br>BIC = 120.52 | Subtask:<br>$F_{(2,250)}$ = 8.875, p < 0.001<br>Time since last visit:<br>$F_{(3,250)}$ = 17.360, p < 0.001<br>Subtask*elapsed time:<br>$F_{(6,250)}$ = 1.298, p = 0.259 |
| **Spline regression** | $R_{adj}$ = 0.174<br>$F_{(7,94)}$ = 4.038<br>p = 0.001<br>RSE = 0.311<br>MSE = 0.104<br>RMSE = 0.322 | $R_{adj}$ = 0.213<br>$F_{(6,48)}$ = 3.092<br>p = 0.009<br>RSE = 0.326<br>MSE = 0.140<br>RMSE = 0.374 | $R_{adj}$ = 0.251<br>$F_{(7,94)}$ = 5.842<br>p < 0.001<br>RSE = 0.387<br>MSE = 0.165<br>RMSE = 0.406 | Subtask:<br>$F_{(2,238)}$ = 9.287, p < 0.001<br>Time since last visit:<br>$F_{(7,238)}$ = 9.015, p < 0.001<br>Subtask*elapsed time:<br>$F_{(14,238)}$ = 1.653, p = 0.066 |

| | | | | |
|---|---|---|---|---|
| | Knots = 180, 2000, 4000, 7500 | Knots = 180, 1000, 2000, 6500 | Knots = 180, 1000, 4000, 6000 | Knots = 180, 2000, 4000, 6000 |
| | AIC = 60.99 | AIC = 42.19 | AIC = 105.48 | |
| | BIC = 84.61 | BIC = 60.26 | BIC = 129.10 | |
| **Generalized additive model** (local smoothing regression of the predictor variable) | $R_{adj}$ = 0.155<br>$F_{(2.57)}$ = 5.831<br>p < 0.001<br>MSE = 0.116<br>Deviance expl.: 17.7%<br>GCV = 0.106<br>AIC = 53.15<br>BIC = 65.69 | $R_{adj}$ = 0.148<br>$F_{(1.99)}$ = 4.059<br>p = 0.017<br>MSE = 0.135<br>Deviance expl.: 17.9%<br>GCV = 0.121<br>AIC = 42.13<br>BIC = 50.14 | $R_{adj}$ = 0.198<br>$F_{(2.6)}$ = 8.037<br>p < 0.001<br>MSE = 0.135<br>Deviance expl.: 21.9%<br>GCV = 0.166<br>AIC = 108.33<br>BIC = 120.41 | Subtask:<br>$F_{(2,253)}$ = 8.533, p < 0.001<br>Time since last visit:<br>$F_{(2,253)}$ = 9.004, p < 0.001 |

In the pointing tasks and the target placement task, the z-score was based on a metric deviation from the target, so that negative correlations were associated with increased memory performance. The blue colour indicates positive correlations in graded weighting. All correlations were Spearman's correlation. Comparison of correlations were conducted with Fishers z-transformation. All models are presented with their key data to explain the variance of spatial memory performance.

## 1.7

**Table 6.** Table of best fitting model formulas and calculations spanning three decades.

| | Direction targets (video) | Direction targets (arrow) | Target placement |
|---|---|---|---|
| Mean and upper to lower standard deviation of zoo-naïve controls (positively transformed (+2) and logarithmized z-scores). | 0.69 (0.599 - 0.779 [log(0.18+2]) | 0.69 (0.604 - 0.775 [log(0.17+2)]) | 0.69 (0.565 - 0.81 [log(0.24+2)]) |
| Best fitting and interpretable model with coefficients | **Power regression**<br>**Intercept:** 0.24306<br>**Log(days):** 0.04114<br>Results are given on log-scale not response scale | Power regression<br>**Intercept:** 0.06619<br>**Log(days):** 0.06495<br>Results are given on log-scale not response scale | Power regression<br>**Intercept:** -0.24826<br>**Log(days):** 0.0930<br>Results are given on log-scale not response scale |
| Timepoint when visitors and controls perform similarly | Mean: 0.24306 + 0.04114*log( **52560 days**) = 0.690<br>SD: 0.24306 + 0.04114*log(5658 days) = 0.599 | Mean: 0.06619 + 0.06495*log( **14965 days**) = 0.691<br>SD: 0.06619 + 0.06495*log(3945 days) = 0.604 | Mean: -0.24826 + 0.0930*log(**24090 days**) = 0.690<br>SD: -0.24826 + 0.0930*log(6300 days) = 0.565 |
| **Timepoint** | | | |
| **Quarter year/ (91.2 days)** | 0.429 (0.366 - 0.491) | 0.59 (0.272 - 0.446) | 0.171 (0.084 - 0.258) |
| **Half year (182.5 days)** | 0.457 (0.406 - 0.509) | 0.404 (0.333 - 0.476) | 0.236 (0.164 - 0.308) |

| | | | |
|---|---|---|---|
| **Year 1 (365 days)** | 0.486 (0.441 – 0.530) | 0.449 (0.390 - 0.509) | 0.300 (0.239 - 0.362) |
| **Year 2 (730 days)** | 0.514 (0.473 – 0.555) | 0.494 (0.440 - 0.549) | 0.365 (0.307 - 0.422) |
| **Year 3 (1095 days)** | 0.531 (0.489 – **0.573**) | 0.521 (0.466 - 0.575) | 0.402 (0.344 - 0.461) |
| **Year 4 (1460 days)** | 0.543 (0.499 – 0.587) | 0.539 (0.483 - **0.596**) | 0.429 (0.368 - 0.490) |
| **Year 5 (1825 days)** | 0.552 (0.506 – **0.598**) | 0.554 (0.495 - 0.613) | 0.450 (0.386 - 0.514) |
| **Year 6 (2190 days)** | 0.559 (0.512 – 0.607) | 0.566 (0.504 - 0.627) | 0.467 (0.400 - 0.534) |
| **Year 7 (2555 days)** | 0.566 (0.516 – 0.616) | 0.576 (0.512 - 0.640) | 0.481 (0.412 - 0.550) |
| **Year 8 (2920 days)** | 0.571 (0.520 – 0.623) | 0.585 (0.518 - 0.651) | 0.493 (0.422 - **0.565**) |
| **Year 9 (3285 days)** | 0.576 (0.523 – 0.629) | 0.592 (0.523 - 0.661) | 0.504 (0.430 - 0.579) |
| **Year 10 (3650 days)** | 0.581 (0.526 – 0.635) | **0.599** (0.528 - 0.670) | 0.514 (0.438 - 0.591) |
| **Year 11 (4015 days)** | 0.584 (0.528 – 0.641) | 0.605 (0.532 - 0.678) | 0.523 (0.445 - 0.601) |
| **Year 12 (4380 days)** | 0.588 (0.531 – 0.646) | 0.611 (0.536 - 0.685) | 0.531 (0.451 - 0.611) |
| **Year 13 (4745 days)** | 0.591 (0.533 – 0.650) | 0.616 (0.540 - 0.692) | 0.539 (0.457 - 0.621) |
| **Year 14 (5110 days)** | 0.594 (0.534 – 0.654) | 0.621 (0.543 - 0.699) | 0.545 (0.462 - 0.629) |
| **Year 15 (5475 days)** | **0.597** (0.536 – 0.658) | 0.625 (0.546 - 0.705) | 0.552 (0.467 - 0.637) |
| **Year 16 (5840 days)** | 0.600 (0.538 – 0.662) | 0.630 (0.549 - 0.710) | 0.558 (0.471 - 0.645) |
| **Year 17 (6205 days)** | 0.602 (0.539 - 0.666) | 0.633 (0.551 - 0.716) | **0.564** (0.475 - 0.652) |
| **Year 18 (6570 days)** | 0.605 (0.540 - 0.669) | 0.637 (0.553 – 0.721) | 0.569 (0.479 - 0.658) |
| **Year 19 (6935 days)** | 0.607 (0.542 – 0.672) | 0.641 (0.556 - 0.726) | 0.574 (0.483 - 0.665) |
| **Year 20 (7300 days)** | 0.609 (0.543 – 0.675) | 0.644 (0.558 - 0.730) | 0.579 (0.486 - 0.671) |
| **Year 21 (7665 days)** | 0.611 (0.544 – 0.678) | 0.647 (0.560 - 0.734) | 0.583 (0.490 - 0.677) |
| **Year 22 (8030 days)** | 0.613 (0.545 – 0.681) | 0.650 (0.562 - 0.739) | 0.588 (0.493 - 0.682) |
| **Year 23 (8395 days)** | 0.615 (0.546 – 0.683) | 0.653 (0.563 - 0.743) | 0.592 (0.496 - 0.687) |
| **Year 24 (8760 days)** | 0.617 (0.547 – 0.686) | 0.656 (0.565 - 0.746) | 0.596 (0.499 – 0.692) |
| **Year 25 (9125 days)** | 0.618 (0.548 – 0.688) | 0.656 (0.567 - 0.750) | 0.599 (0.502 – 0.697) |
| **Year 26 (9490 days)** | 0.620 (0.549 – 0.691) | 0.661 (0.568 - 0.754) | 0.603 (0.504 – 0.702) |
| **Year 27 (9855 days)** | 0.621 (0.550 – 0.693) | 0.664 (0.570 - 0.757) | 0.607 (0.507 – 0.706) |

| | | | |
|---|---|---|---|
| **Year 28 (10220 days)** | 0.623 (0.551 – 0.695) | 0.666 (0.571 – 0.760) | 0.610 (0.509 – 0.711) |
| **Year 29 (10585 days)** | 0.624 (0.551 – 0.697) | 0.668 (0.573 - 0.763) | 0.613 (0.512 – 0.715) |
| **Year 30 (10950 days)** | 0.625 (0.552 – 0.699) | 0.670 (0.574 - 0.767) | 0.616 (0.514 – 0.719) |
| ### | | | |
